# Genome-wide definition of the CosR regulon and DNA-binding properties in *Campylobacter jejuni*

**DOI:** 10.64898/2026.08.04.742711

**Authors:** Elena Chiti, Tobia Odorici, Mateusz Noszka, Jakub Muraszko, Annamaria Zannoni, Anna Zawilak-Pawlik, Andrea Vannini, Davide Roncarati

**Affiliations:** Department of Pharmacy and Biotechnology, University of Bologna, Via Selmi 3, 40126 Bologna, Italy; Department of Microbiology, Hirszfeld Institute of Immunology and Experimental Therapy, Polish Academy of Sciences, Weigla 12, 53-114 Wroclaw, Poland; University of Würzburg, Institute of Molecular Infection Biology, Department of Molecular Infection Biology II, Würzburg, Germany

**Keywords:** *Campylobacter jejuni*, bacterial transcriptional regulation, ChIP-seq, protein-DNA interactions, CosR, oxidative stress response, genome-wide regulon

## Abstract

CosR is an essential OmpR-family transcriptional regulator of *Campylobacter jejuni*, but its direct regulon and DNA-binding properties in vivo remain poorly defined. Here, we used ChIP-seq with a functional chromosomal CosR::3×FLAG allele to define the genome-wide CosR-binding landscape. CosR binding was strongly enriched at promoters and accumulated around transcription start sites, consistent with a primary role in transcriptional control. Functional analysis of promoter-bound targets revealed significant enrichment for translation- and transcription-related functions, identifying CosR as a regulator of core cellular processes. Motif analysis of summit-centered ChIP-seq regions identified a CosR-associated bipartite sequence signature characterized by TTAA-like elements separated by an A/T-rich spacer. DNase I footprinting confirmed direct promoter binding at nucleotide resolution and revealed heterogeneous architectures, including single, multipartite, and bidirectional binding arrangements. A footprint-derived motif was significantly similar to the ChIP-derived motif, supporting a shared recognition signature across in vivo-enriched regions and in vitro-protected segments. Footprinting also validated binding at non-coding RNA promoters and at the *cosR* promoter, indicating autoregulation.

Hydrogen peroxide treatment differentially remodeled CosR promoter occupancy in vivo, reducing binding at translation-associated promoters while increasing enrichment at other targets. Redox-dependent footprinting showed that oxidative conditions directly impaired CosR binding at selected promoters, consistent with previously reported C218-dependent redox modulation. Despite these opposite occupancy patterns, most tested transcripts decreased after oxidative stress, indicating that CosR redox responsiveness is integrated with broader stress-dependent regulatory inputs. Together, these data define CosR as a condition-responsive regulator linking promoter recognition, core physiology and oxidative-stress-associated transcriptional remodeling.

**IMPORTANCE:** *Campylobacter jejuni* is a major foodborne pathogen that must adapt to changing oxygen levels and stress conditions during transmission and infection. CosR is essential for this bacterium, but previous studies could not clearly distinguish direct regulation from secondary effects caused by perturbing an essential protein. By mapping CosR binding across the genome and validating binding at nucleotide resolution, this study shows that CosR directly targets promoters associated with fundamental cellular functions, particularly translation and RNA metabolism, showing that the role of CosR extends well beyond oxidative-stress regulation. We also show that a brief oxidative challenge redistributes CosR occupancy among promoters in a target-dependent manner. These findings identify CosR as a condition-responsive regulator linking the gene-expression machinery and other core physiological functions to stress adaptation, and provide a framework for understanding how *C. jejuni* adjusts its physiology in changing environments.

## INTRODUCTION

*C. jejuni* is a Gram-negative microaerophilic bacterium and a major foodborne pathogen worldwide (1–3). *Campylobacter* spp. are prevalent in poultry and other livestock, but also in pets, including cats and dogs, and *C. jejuni* can survive outside the animal host and colonize the human gastrointestinal tract (4). Human infection is mainly associated with the ingestion of contaminated food, particularly raw or undercooked meat, raw milk, or untreated water (5, 6), and can cause campylobacteriosis, a foodborne illness characterized by diarrhea, abdominal pain, and fever (2, 3, 6). Although usually self-limiting, infection can become severe in vulnerable individuals and lead to complications such as bacteremia, reactive arthritis, or Guillain-Barré syndrome (2, 3). In severe cases, antibiotic treatment may be required (3), but multidrug-resistant *Campylobacter* contributes to the broader global concern over bacterial antimicrobial resistance (7). As a microaerophilic organism inhabiting diverse environmental niches, *C. jejuni* is frequently exposed to highly variable environmental conditions and multiple stress factors (8–10). Rapid transcriptional reprogramming is therefore central to its adaptation and relies on a compact but complex regulatory network (4).

CosR is an essential transcriptional regulator belonging to the OmpR/PhoB family of response regulators, whose homologues are widespread in Campylobacteria, including *Helicobacter* and *Wolinella* (11). In *C. jejuni*, CosR is considered an orphan response regulator, as it lacks a cognate sensor kinase and the conserved phosphate-accepting aspartate is absent. CosR is therefore thought to operate as an autonomous transcriptional regulator via a phosphorylation-independent mechanism (12).

Recent structural studies have provided important insight into CosR DNA binding. Cryo-EM and X-ray crystallography revealed that CosR forms a dimer with an N-terminal receiver domain connected by a flexible linker to a C-terminal winged helix-turn-helix DNA-binding domain (13). Few CosR binding sites have been identified, preventing consensus sequence definition. However, crystallographic structures of CosR complexed with a 21-bp DNA fragment from the *cmeABC* promoter, previously identified as a CosR-protected region by footprinting, showed that DNA binding imposes a switch from a symmetric to an asymmetric arrangement of the CosR dimer, in which the DNA-binding domains engage the target DNA through major- and minor-groove contacts (13). One protomer adopts a compact conformation with strong interactions between the N- and C-terminal domains, whereas the other remains in an extended conformation. These structural features resemble those of the *H. pylori* HP1043/HsrA regulator, consistent with the near-perfect amino acid identity in their DNA-binding domains (14). The observed conformational flexibility may also provide a structural basis for coupling DNA recognition to yet unidentified environmental signals, possibly including oxidative stress, as suggested by the redox sensitivity of the conserved C218 residue (13, 15).

The essentiality of *cosR* has hampered detailed definition of its in vivo regulatory role. Initial studies based on antisense-mediated *cosR* knockdown and proteomic analysis identified 32 CosR-responsive proteins involved in processes including macromolecule biosynthesis, metabolism, gene regulation, and oxidative stress response (11). Subsequent transcriptomic analysis by DNA microarrays under antisense-mediated *cosR* knockdown conditions identified 93 differentially expressed genes linked to drug and ROS resistance, motility, and functions important for *C. jejuni* viability (16). These studies suggested that CosR acts as a broad regulator, but their interpretation is complicated by the need to perturb an essential protein. The previously used 8-h PNA-mediated CosR knockdown may severely affect cell physiology and generate pleiotropic secondary responses, in addition to possible PNA-associated off-target effects or toxicity. Constitutive overexpression poses a related problem, as CosR levels are altered throughout growth and may reflect long-term adaptation rather than direct regulatory effects. Thus, many CosR-responsive changes detected under these conditions may represent indirect consequences of cellular imbalance rather than direct promoter control. Consistently, proteomic and transcriptomic screens identified only partially overlapping CosR-responsive candidates, likely reflecting both the different temporal and regulatory layers captured by mRNA and protein-level analyses and the accumulation of secondary physiological effects. Moreover, direct CosR binding has been validated for only a small number of promoters (11, 16, 17).

We performed genome-wide CosR ChIP-seq in *C. jejuni* using a functional chromosomal CosR::3×FLAG strain. We mapped 208 high-confidence CosR-binding sites, integrated the ChIP-seq dataset with transcription start site-aware promoter annotation, and validated selected promoters by ChIP-qPCR, EMSA, and footprinting. Furthermore, we investigated oxidative stress-dependent remodeling of CosR promoter occupancy by ChIP-qPCR, redox-dependent footprinting, and transcriptional profiling. Our results define CosR as a central regulator of core cellular processes, including translation, transcription, energy metabolism, and stress-associated functions, and reveal dynamic, target-specific modulation of CosR-DNA binding under oxidative conditions.

## MATERIALS AND METHODS

Detailed experimental and computational procedures are provided in Supplementary Materials and Methods.

### Bacterial strains and CosR::3×FLAG construction

*C. jejuni* NCTC11168 wild-type and CosR::3×FLAG strains were grown under microaerobic conditions at 42°C in BHI or Columbia medium. The chromosomal CosR::3×FLAG strain was generated by natural transformation and homologous recombination using a pUC18-derived construct carrying C-terminally 3×FLAG-tagged cosR and a kanamycin-resistance cassette. Correct integration was verified by PCR and Sanger sequencing. Strains, plasmids, and oligonucleotides are listed in Supplementary Tables S1 and S2.

### ChIP, expression analysis, and DNA-binding assays

For ChIP-seq, wild-type and CosR::3×FLAG cultures were crosslinked immediately upon reaching OD_600_ 1.2. Chromatin was sheared, immunoprecipitated with anti-FLAG magnetic agarose, de-crosslinked, purified, and subjected to paired-end sequencing. ChIP-qPCR was normalized to an intragenic *gyrA* region lacking detectable CosR enrichment. For oxidative-stress assays, cultures were exposed to 1.5 mM H_2_O_2_ for 20 min before ChIP or RNA extraction. RT-qPCR data were calculated by the ΔΔCt method using 16S rRNA as internal reference. CosR and CosR::3×FLAG accumulation was assessed by western blotting, and growth was monitored in BHI broth. Recombinant C-terminally STREP-tagged CosR was purified from *E. coli* BL21(DE3) and used for EMSA and DNase I footprinting, including redox-dependent assays performed with DTT or H_2_O_2_.

### ChIP-seq processing, annotation, motif analysis, and statistics

Reads were processed in Galaxy, aligned to the *C. jejuni* NCTC11168 genome with Bowtie2, and analyzed with MACS2 and irreproducible discovery rate analysis. Downstream analyses used WT-normalized CosR peak summits with FC ≥ 1.5; the stringent core subset was defined by FC ≥ 2.5. TSS assignment was based on the published SuperGenome annotation (18) supplemented by manually curated TSSs. Functional classification used eggNOG annotation followed by manual curation. Motif discovery was performed with MEME-ChIP, followed by PWM refinement and comparison with footprint-derived motifs. Statistical analyses used Student’s t test or Fisher’s exact test, as appropriate (18).

### Data availability

ChIP-seq data will be deposited in a public repository, with the accession number provided during revision.

## RESULTS

### Genome-wide identification of in vivo CosR-enriched regions in Campylobacter jejuni

To define the in vivo DNA-binding landscape of CosR in *C. jejuni*, we initially attempted ChIP using a polyclonal anti-CosR antibody, but this approach did not provide sufficient enrichment and was not pursued further. We therefore generated a strain expressing a functional C-terminally 3×FLAG-tagged CosR and assessed whether the tag affected CosR accumulation or bacterial growth. Western blot analysis with the polyclonal anti-CosR antibody showed comparable CosR and CosR::3×FLAG levels in wild-type and tagged strains across different growth phases, with the expected mobility shift caused by the FLAG tag (Fig. S1A). Anti-FLAG immunoblotting confirmed specific detection of the tagged protein only in the CosR::3×FLAG strain (Fig. S1B). In addition, the tagged strain showed growth kinetics and generation time comparable to the wild type (Fig. S1C,D). These data support the use of the CosR::3×FLAG strain for ChIP-seq analysis. ChIP-seq was performed on three independent biological replicates of the CosR::3×FLAG strain, with the wild-type strain processed in parallel as a negative control.

Across all samples, 7.1–8.4 million paired-end reads were generated, with >7 million paired reads retained after trimming. After mapping, independent peak calling, and reciprocal intersection of pairwise IDR-supported peak sets, the uniquely mapped dataset yielded 268 reproducible binding regions (Fig. 1A).

**Figure 1.**
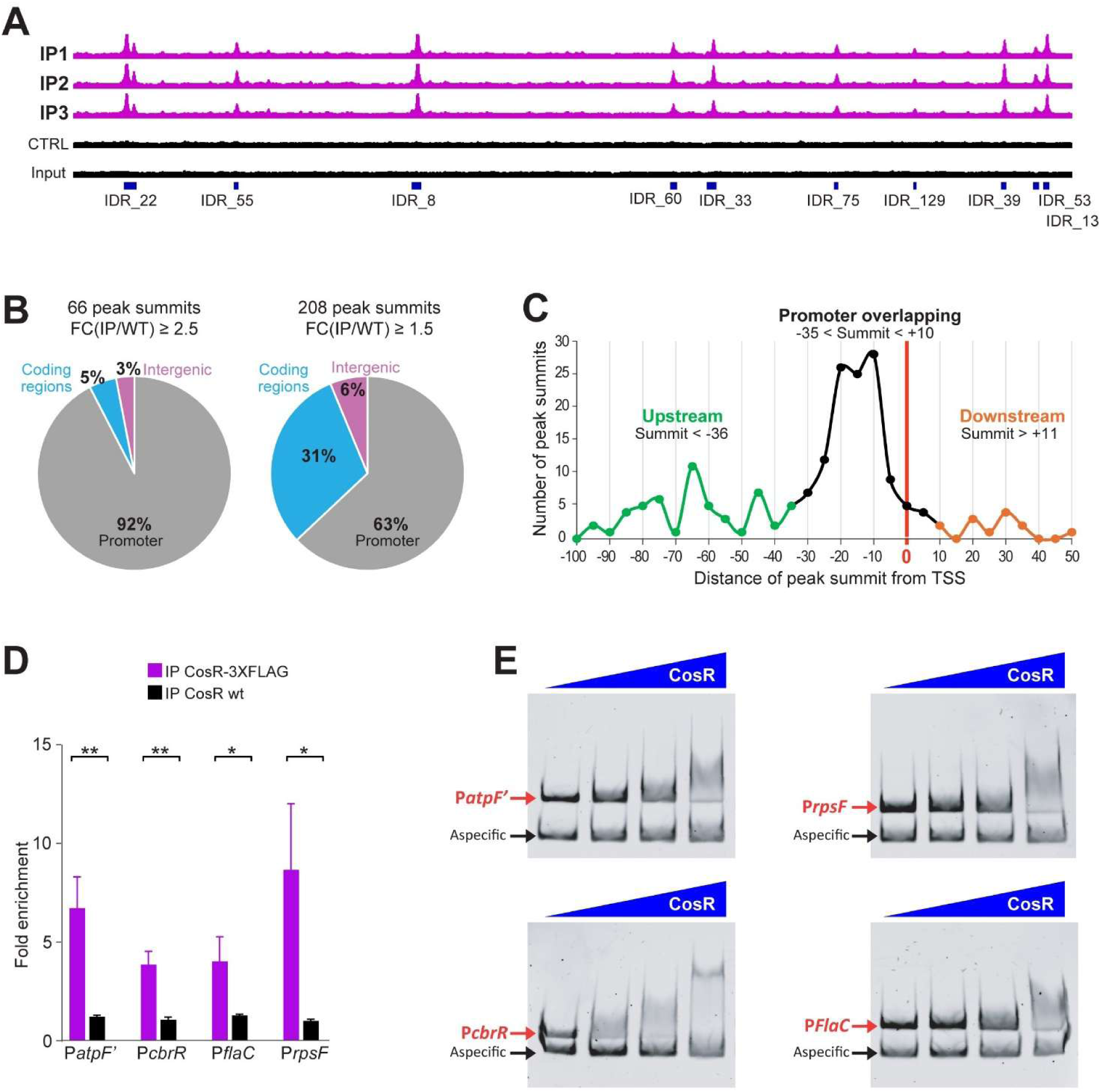
Genome-wide identification and validation of CosR-bound promoter regions in *Campylobacter jejuni*. (A) Representative 100-kb genomic window showing ChIP-seq coverage profiles from the three biological replicates of the CosR::3×FLAG strain (IP1–IP3), together with the wild-type mock-IP control (CTRL) and the input DNA track. Blue boxes indicate representative irreproducible discovery rate (IDR)-supported CosR-enriched regions. (B) Genomic classification of WT-filtered CosR peak summits in the stringent core subset (66 summits, FC [IP/WT] ≥ 2.5) and in the extended dataset (208 summits, FC [IP/WT] ≥ 1.5). Summits were classified as promoter-associated, coding region, or intergenic. (C) Distribution of promoter-associated summit positions relative to the transcription start site (TSS). Based on summit-to-TSS distance, regulatory units were classified as promoter-overlapping (−35 to +10; black), upstream (−100 to −36; green), or downstream (+11 to +50; orange). (D) ChIP-qPCR validation of selected promoter-associated CosR targets. Enrichment of the indicated promoter regions (*atpF’*, *cbrR*, *flaC*, *rpsF*) was measured in independent ChIP experiments using the CosR::3×FLAG strain (magenta) and the wild-type strain (black). Values are shown as fold enrichment relative to the corresponding input DNA. Error bars indicate standard deviation from four biological replicates. Statistical significance was assessed by Student’s t test. (E) EMSA validation of the same promoter targets shown in panel D. Increasing concentrations of recombinant CosR (0, 0.1, 0.2, or 0.4 µg corresponding to 0, 186, 370, 740 nM) were incubated with promoter probes derived from P*atpF’*, P*rpsF*, P*cbrR*, and P*flaC*. All four probes formed specific shifted complexes in the presence of CosR. The non-specific control probe (aspecific) mapping on the *recJ* CDS is indicated. *, P < 0.05; **, P < 0.01; ***, P < 0.001

Because exclusion of multimapping reads distorted peak profiles at multicopy loci, we performed a parallel multimapping-aware analysis. Multimapping-aware analysis corrected summit positions at rRNA promoters and recovered nine additional reproducible peaks. The final dataset comprised 277 reproducible peak regions. Manual inspection identified 13 regions containing more than one local maximum, resulting in 290 candidate summits overall (Supplementary Table S3). WT-normalized enrichment was then quantified in summit-centered windows (±50 nt), and downstream analyses were restricted to 208 summits with FC ≥ 1.5, including a more stringent “core” subset of 66 summits with FC ≥ 2.5.

### Genomic distribution and promoter positioning of CosR-binding sites

Annotation of the 208 WT-filtered summits revealed a strong promoter bias among higher-confidence CosR-binding sites (Supplementary Table S4). Promoter assignments were based on the curated primary-TSS dataset described in the Materials and Methods section, comprising SuperGenome-derived NCTC11168 primary TSSs supplemented with 33 manually refined promoter assignments (Supplementary Table S5). In the stringent core subset (66 summits with FC ≥ 2.5), 61 sites (92%) mapped to promoters, whereas 3 fell within coding regions and 2 within intergenic regions (Fig. 1B). Expanding the analysis to all 208 summits with FC ≥ 1.5 reduced the promoter-associated fraction to 63% (131 sites), with 64 summits mapping within coding regions and 13 within intergenic regions (Fig. 1B). Because TSS usage in *C. jejuni* can vary across growth and stress conditions, some coding-region or intergenic assignments may reflect condition-dependent or incompletely annotated promoters rather than promoter-independent CosR binding. The 131 promoter-associated summits were then linked to 182 regulatory units, which were used for downstream positional analyses.

The positional distribution of these regulatory units relative to the TSS revealed a marked accumulation of summit positions in the core promoter region (Fig. 1C). Based on the frequency distribution of summit-to-TSS distances, regulatory units were classified into three positional groups: promoter-overlapping (−35 to +10), upstream (−100 to −36), and downstream (+11 to +50). Among the 182 promoter-associated regulatory units, 120 fell within the promoter-overlapping class, 51 were located upstream, and 11 downstream of the TSS (Supplementary Table S6).

### Validation of selected CosR targets by ChIP-qPCR and EMSA

To validate the ChIP-seq dataset, we selected four high-confidence promoter-associated targets for further analysis: *atpF′*, *rpsF*, *cbrR*, and *flaC*. These genes play important roles in *C. jejuni* physiology and pathogenesis: *atpF′*, an ATP synthase subunit required for energy metabolism and survival under changing osmotic conditions (19); *rpsF*, encoding the 30S ribosomal protein S6 essential for translation; *flaC*, a secreted flagellin involved in motility and host interactions (20); and *cbrR*, a regulator linking bile resistance with motility and biofilm formation (21, 22). These loci were chosen based on strong enrichment and clear promoter localization in the ChIP-seq profiles. Enrichment of the corresponding promoter regions was first assessed by ChIP-qPCR on independent biological material. In all cases, the CosR::3×FLAG strain showed significantly higher recovery than the wild-type control, confirming specific enrichment of CosR-bound promoter DNA *in vivo* (Fig. 1D). Depending on the target, enrichment ranged from approximately 4-to 9-fold over background.

We next tested whether CosR could bind these regions directly *in vitro*. EMSA assays performed with recombinant CosR and promoter probes corresponding to the same four loci showed specific shifted complexes for all targets (Fig. 1E). Although the apparent binding strength varied among promoters, all four probes were bound by CosR, whereas the non-specific control probe derived from the *recJ* CDS did not show detectable binding.

### Functional enrichment of genes associated with promoter-bound CosR targets

To explore the functional organization of the CosR regulon, we reconstructed operons associated with the core promoter-bound summits using the curated primary-TSS dataset described in Materials and Methods. All genes within these operons were then assigned to COG functional categories, using eggNOG-based annotation (Supplementary Table S7) followed by manual curation of unresolved cases (Supplementary Table S8). Compared with the overall genomic background (Fig. S1E), genes associated with CosR-bound promoter regions were significantly enriched in COG J (translation, ribosomal structure and biogenesis, including rRNA- and tRNA-associated targets) and COG K (transcription) (Fig. 2A). When promoter-associated targets were subdivided according to summit position relative to the TSS, distinct functional biases emerged. The promoter-overlapping group showed selective enrichment of COG J and COG D (cell cycle control, cell division, chromosome partitioning), whereas the upstream group was enriched in COG K and COG N (cell motility) (Fig. 2B). These data indicate that CosR binding is not only concentrated at promoter regions but is also functionally structured according to its positional relationship with the TSS.

**Figure 2.**
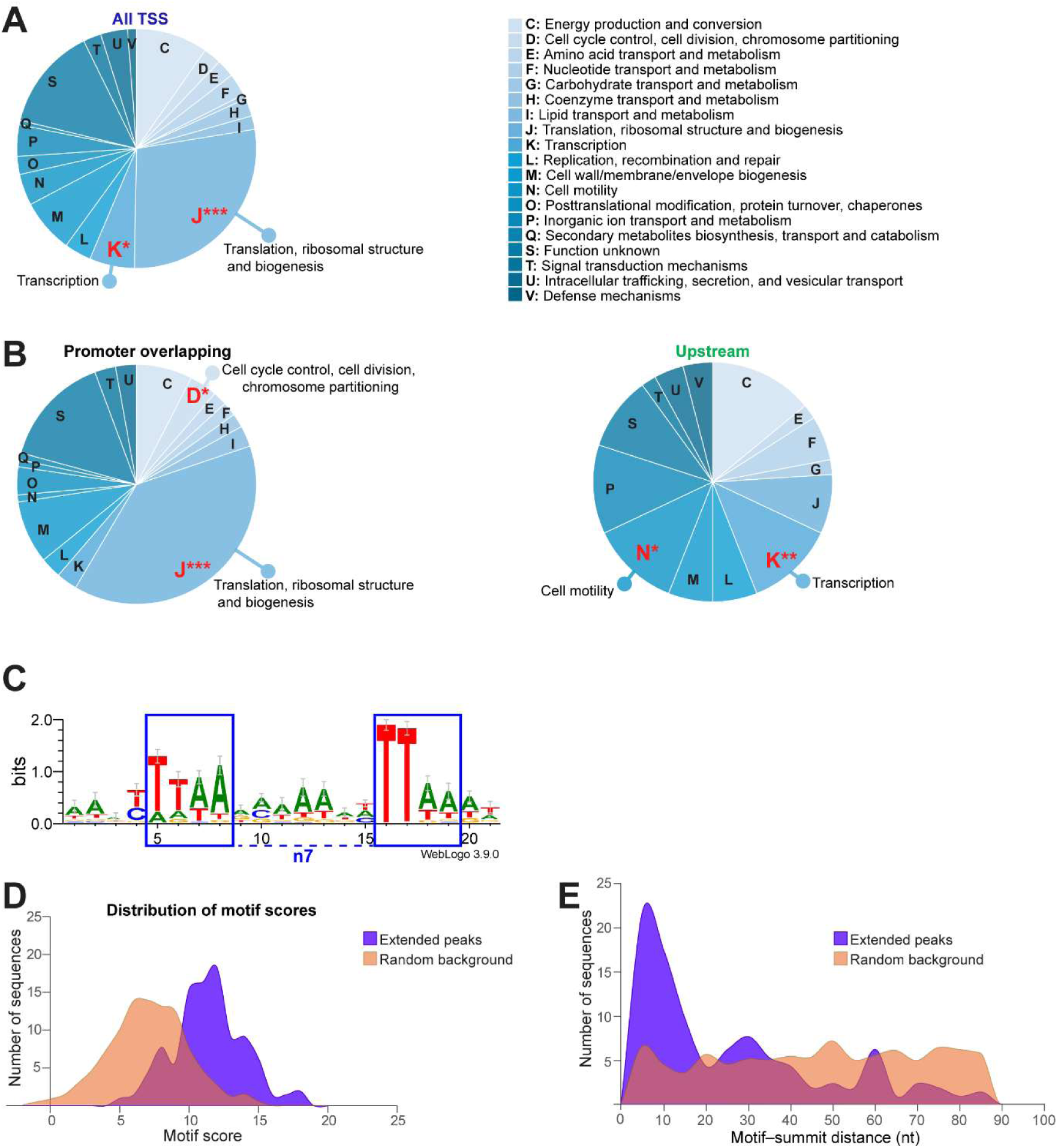
Functional enrichment and peak-centered motif analysis of promoter-associated CosR targets. (A) COG classification of genes associated with promoter-bound CosR targets. Significantly enriched categories relative to the genomic background are highlighted in red; enrichment was assessed by Fisher’s exact test. (B) COG classification of genes associated with promoter-overlapping and upstream CosR-bound regulatory units. Significantly enriched categories relative to the genomic background are highlighted in red; enrichment was assessed by Fisher’s exact test. (C) Sequence logo derived from de novo motif discovery on ±100-bp regions centered on the 208 WT-filtered CosR ChIP-seq summits, showing the candidate ChIP-derived CosR-associated motif. (D) Distribution of motif scores obtained by rescanning the 208 WT-filtered summit-centered CosR-bound sequences with the refined ChIP-derived PWM. Purple indicates extended CosR peaks; orange indicates 1,040 peak-excluded random genomic background sequences sampled at five-fold the size of the extended peak set. (E) Distribution of distances between motif centers and sequence centers. For extended CosR peaks, the sequence center corresponds to the ChIP-seq summit; for random background sequences, it corresponds to the center of the sampled genomic interval.

### Peak-centered motif analysis identifies a candidate CosR-associated sequence signature

To identify sequence features associated with CosR-bound regions, we performed de novo motif discovery on summit-centered ChIP-seq sequences. Initial analysis of the stringent core set of 66 summits yielded a motif closely resembling the canonical bacterial −10 promoter element (Fig. S2A). Because more than 90% of core summits mapped to promoter regions, this signal likely reflected the strong promoter bias of the core dataset rather than a CosR-specific binding determinant and was therefore not considered further.

We next analyzed the extended set of 208 WT-filtered summit-centered sequences using 201-bp regions centered on each summit. This analysis identified a distinct bipartite motif, approximated by TTAA-n7-TTAA (Fig. 2C), characterized by TTAA-like elements separated by an A/T-rich spacer. To evaluate its enrichment and positional distribution, summit-centered peak sequences were rescanned with the refined PWM and compared with peak-excluded random genomic background sequences sampled at five-fold the size of the extended peak set. CosR-bound regions showed a clear shift toward higher motif scores compared with the random background (Fig. 2D), indicating that this motif is enriched within ChIP-seq peak regions. In addition, motif centers were strongly concentrated close to ChIP-seq summits, whereas motif positions in random genomic regions showed a broader, near-uniform distance distribution relative to the center of the sampled genomic interval (Fig. 2E).

As an internal consistency check, rescanning of the stringent core subset with the extended-set PWM showed motif-score and motif-to-summit distance distributions comparable to those of the extended dataset (Supplementary Table S9), supporting recovery of the ChIP-derived motif also among high-confidence CosR peaks (Fig. S2B, C). Together, these analyses support the enrichment and summit-proximal positioning of this AT-rich sequence signature within CosR-bound regions, supporting its use as a candidate ChIP-derived CosR-associated motif.

### High-resolution mapping of CosR-protected regions supports the ChIP-seq-defined binding landscape

To define CosR-binding sites at nucleotide resolution, we performed DNase I footprinting on a representative set of promoter regions selected from the CosR ChIP-seq dataset. In addition to the *cbrR* and *rpsF* promoters, previously validated by ChIP-qPCR and EMSA, we analyzed *hydA*, which contributes to hydrogen utilization and bacterial fitness during host colonization, and the bidirectional promoter shared by the divergent *cj0459c* and *nusA* transcription units, the latter encoding the essential transcription elongation factor NusA (Fig. 3A). In all four cases, footprinting confirmed direct CosR binding within promoter-proximal regions. However, the organization of the protected segments differed among targets. While P*rpsF* showed a single protected region, P*cbrR* and P*hydA* displayed multiple protected segments, consistent with the presence of more than one CosR-binding site within the same promoter region. In the *cj0459c/nusA* bidirectional promoter, a single protected region was compatible with promoter association for both divergent transcription units.

**Figure 3.**
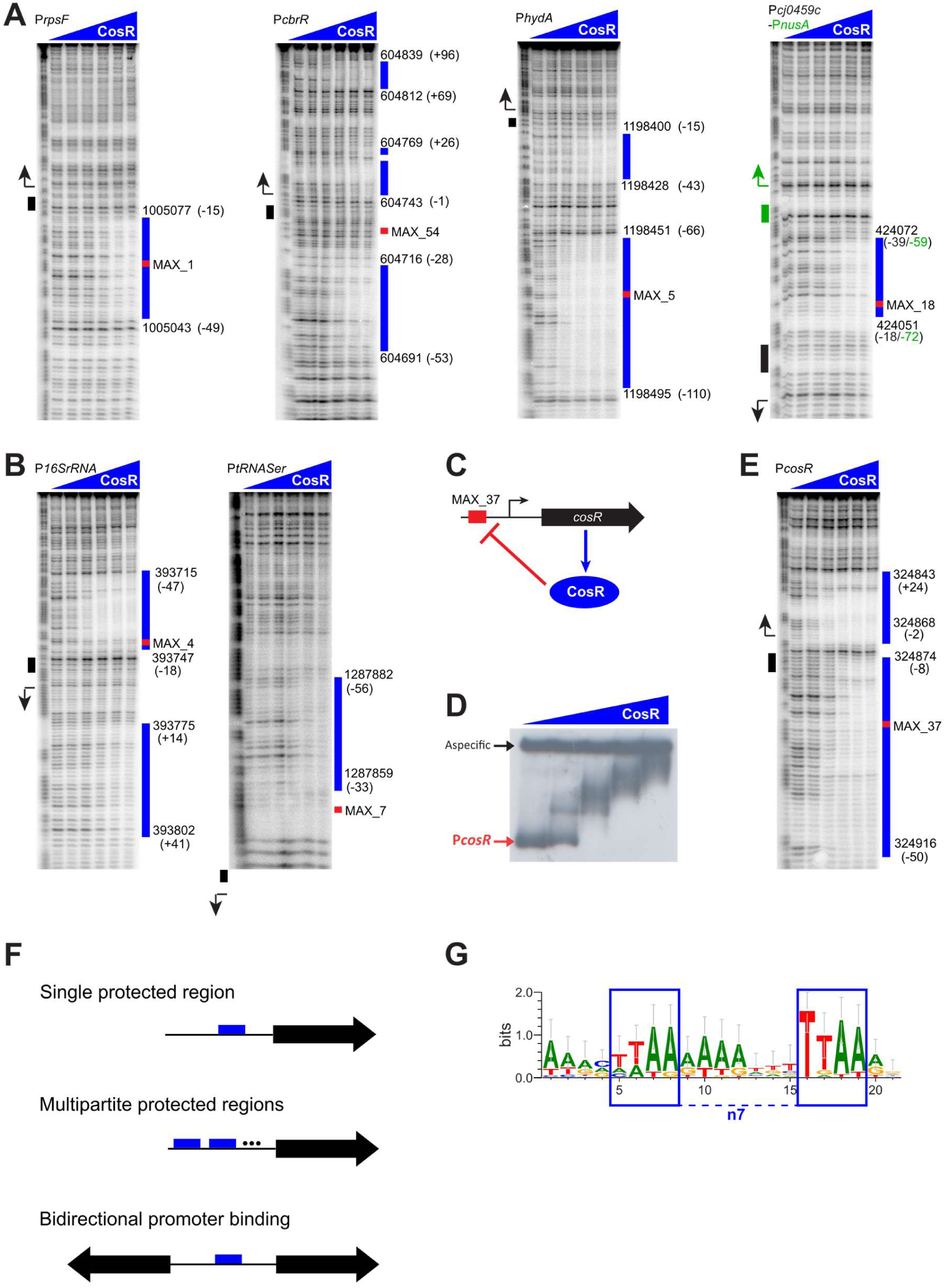
DNase I footprinting defines direct CosR-bound promoter regions and supports autoregulation. (A) DNase I footprinting of representative protein-coding promoters bound by CosR. The validated targets P*cbrR* and P*rpsF*, the newly identified target P*hydA*, and the bidirectional P*cj0459c*/P*nusA* promoter region are shown. Blue bars indicate protected regions, red boxes indicate the corresponding ChIP-seq summit, and numbers in parentheses indicate the position relative to the associated TSS. (B) DNase I footprinting of representative non-coding RNA promoters, including P*16SrRNA* and P*tRNASer*. (C) Model of CosR autoregulation based on the presence of summit MAX_37 in the *cosR* promoter region. (D) EMSA showing direct binding of CosR to the *cosR* promoter. (E) DNase I footprinting of the *cosR* promoter, confirming direct promoter binding by CosR. (F) Schematic summary of the three recurrent CosR promoter-binding architectures identified by DNase I footprinting: promoters containing a single protected region, promoters displaying multipartite binding characterized by two or more protected regions, and bidirectional promoters in which a single protected region is shared between divergent transcription units. Schematics are conceptual and are not intended to represent the exact number, spacing, or position of protected regions. (G) Sequence logo derived from alignment of the CosR-protected regions identified by DNase I footprinting.

We next extended the analysis to representative non-coding RNA promoters identified by ChIP-seq. Footprinting of 16S rRNA and tRNA-Ser promoter regions again confirmed direct CosR binding, with the rRNA promoter showing a bipartite protection pattern and the tRNA promoter a single protected region (Fig. 3B). Additional footprinting analyses of other direct targets, including *atpF’*, *rpmJ*, *flaC*, *ogt/acnB*, *atpE*, *rplS*, *cheY*, and *gyrA*, are shown in Fig. S3A-C. Together, these experiments showed that all tested ChIP-seq targets were positive in DNase I footprinting, including both stringent core summits and lower-enrichment extended targets, strongly supporting the overall quality of the CosR ChIP-seq dataset.

Considering all experimentally characterized promoters, including the additional targets shown in Fig. S3, three recurrent CosR-binding architectures emerged (Fig. 3F). Promoters contained either a single protected region, multiple protected regions within the same promoter, or a single protected region associated with a bidirectional promoter shared by divergent transcription units. Although the number and relative arrangement of protected regions varied among individual promoters, these three classes captured the major binding architectures observed across the footprinting dataset.

Comparison of ChIP-seq summits and footprint-defined protected regions further highlighted a strong convergence between the two approaches. In most cases, the summit mapped within a protected region, as observed for *rpsF*, *hydA*, and *nusA*, or between two protected regions, as seen for multipartite targets such as *cbrR* and *acnB*. These patterns indicate that nearby CosR-binding sites are often resolved as a single enriched region by ChIP-seq, whereas DNase I footprinting distinguishes their underlying fine structure. Only a few targets, including *tRNASer*, *cheY*, and *gyrA*, showed a summit outside the protected region, but in all cases the displacement remained limited to the immediate vicinity or within approximately 50 bp.

All DNase I footprinting assays were performed at 42°C, the physiological growth temperature of *C. jejuni* in birds. To assess whether temperature might influence CosR DNA binding, as reported for other transcriptional regulators, the *hydA* promoter was also analyzed at 25°C, the standard temperature commonly used for in vitro footprinting assays. The protection pattern was identical under the two conditions (Fig. S3D), indicating that CosR binding to the *hydA* promoter is not detectably affected by temperature in this assay.

Since autoregulation is common among bacterial transcriptional regulators, the presence of a ChIP-seq summit (MAX_37) within the *cosR* promoter prompted us to test whether CosR binds its own promoter directly (Fig. 3C). Both EMSA (Fig. 3D) and DNase I footprinting (Fig. 3E) confirmed direct binding of CosR to the *cosR* promoter, supporting an autoregulatory loop.

### Comparison of ChIP-derived and footprint-derived CosR binding motifs

Motif analysis of the 23 DNase I footprint-protected regions identified a sequence signature (Fig. 3G) that was significantly similar to the ChIP-derived motif, as assessed by Tomtom comparison (overlap, 20 nt; q-value = 1.3 × 10⁻⁴). The ChIP-derived and footprint-derived motifs converged on a bipartite architecture characterized by two TTAA-like elements separated by approximately 7 nt, together with an A/T-rich central spacer and additional conserved positions in the flanking regions (Fig. 3G). These results support a shared CosR-associated recognition signature across the in vivo ChIP-seq dataset and the in vitro footprint-protected regions. Differences in individual nucleotide preferences likely reflect the different nature of the input sequences, with ChIP-seq using summit-centered enriched regions and footprinting using fewer, experimentally protected segments from structurally heterogeneous promoters.

### Oxidative stress remodels CosR promoter occupancy and target gene expression

Because CosR has been implicated in the oxidative stress response of *C. jejuni*, we asked whether a brief H₂O₂ pulse would alter its promoter occupancy in vivo. CosR::3×FLAG cultures were exposed to hydrogen peroxide for 20 min, and binding to selected promoters was quantified by ChIP-qPCR. Oxidative stress produced two distinct occupancy patterns. CosR enrichment increased at a subset of promoters, including P*atpF′*, P*cbrR*, P*flaC*, and P*cosR*, whereas P*ahpC* remained negative under both conditions (Fig. 4A). By contrast, CosR occupancy decreased at translation-associated promoters, including P*rpsF*, P*rplS*, P*rpmJ*, and P*tRNASer* (Fig. 4B). Thus, oxidative stress does not globally increase or decrease CosR binding, but remodels CosR occupancy in a target-dependent manner.

**Figure 4.**
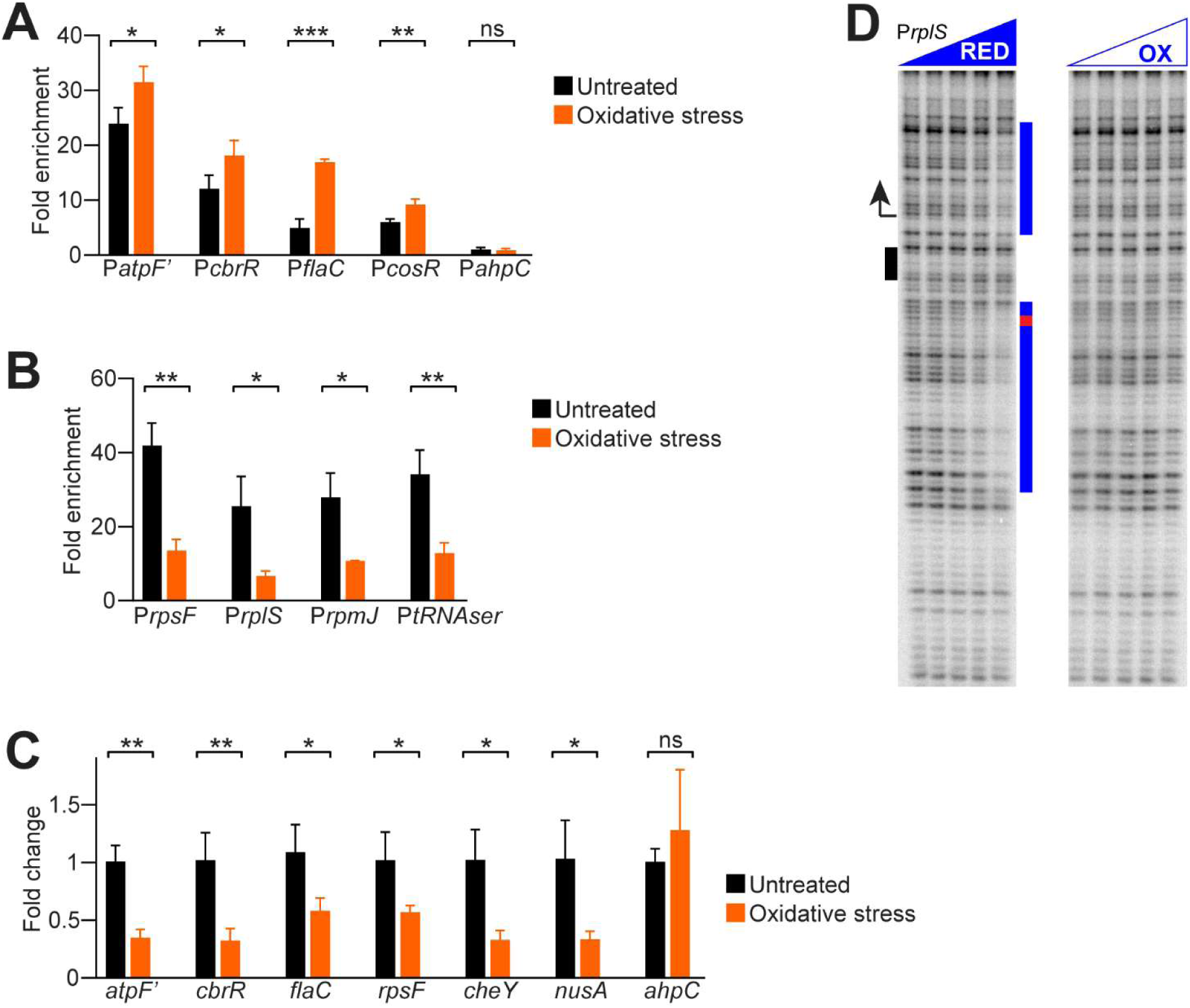
Oxidative stress differentially remodels CosR promoter occupancy and target gene expression. (A) ChIP-qPCR analysis of CosR occupancy at selected promoters showing increased enrichment after a brief oxidative stress treatment. Enrichment was measured in untreated cultures and after 20 min exposure to H₂O₂ for P*atpF’*, P*cbrR*, P*flaC*, P*cosR*, and P*ahpC*. P*ahpC* remained negative under both conditions. (B) ChIP-qPCR analysis of CosR occupancy at translation-associated promoters showing reduced enrichment after oxidative stress, including P*rpsF*, P*rplS*, P*rpmJ*, and P*tRNASer*. (C) RT-qPCR analysis of selected CosR-associated target genes after oxidative stress. Transcript levels of *atpF’*, *cbrR*, *flaC*, *rpsF*, *cheY*, and *nusA* were reduced relative to untreated cultures, whereas *ahpC* showed no significant change. Values are shown as fold change relative to the untreated condition. (D) DNase I footprinting of the *rplS* promoter under reducing and oxidizing conditions. CosR-dependent protection was detected under reducing conditions and was strongly reduced or lost under oxidizing conditions. Blue bars indicate protected regions. Error bars indicate standard deviation from three biological replicates. Statistical significance was assessed by Student’s t test. *, P < 0.05; **, P < 0.01; ***, P < 0.001

To determine whether these occupancy changes were associated with altered transcription, transcript levels of selected targets were measured by RT-qPCR after hydrogen peroxide treatment. Most tested genes showed reduced transcript abundance under oxidative stress, including *atpF′*, *cbrR*, *flaC*, *rpsF*, *cheY*, and *nusA*, whereas *ahpC* showed no significant change (Fig. 4C). Therefore, changes in CosR occupancy were not sufficient to predict transcriptional output under oxidative stress.

We next tested whether oxidation could directly affect CosR-DNA binding in vitro. DNase I footprinting under reducing and oxidizing conditions showed that CosR-dependent protection at the *rplS* promoter was detected under reducing conditions but was strongly reduced or lost in the presence of H₂O₂ (Fig. 4D). A similar reduction in CosR-dependent protection was observed at the *atpE* promoter under oxidizing conditions (Fig. S3E). These data indicate that oxidative conditions can directly impair CosR binding to multiple target promoters. For *rplS*, this is consistent with the reduced in vivo occupancy observed at translation-associated loci.

## DISCUSSION

This study provides the first genome-wide definition of the direct CosR-binding landscape in *C. jejuni*. By combining ChIP-seq with ChIP-qPCR, EMSA, and DNase I footprinting, we show that CosR binding is strongly enriched at promoter regions and frequently positioned close to transcription start sites. This supports a primary role for CosR in transcriptional control and expands the set of direct CosR-associated promoter targets from a handful of previously validated promoters to more than one hundred ChIP-supported promoter-associated sites. Importantly, the functional organization of promoter-bound core targets indicates that CosR is not restricted to oxidative-stress defense but instead controls a broad set of central cellular functions.

Manual functional classification of genes belonging to core promoter-bound operons highlighted translation and RNA metabolism as the most prominent functional axis of the CosR regulon (Fig. 5). This group includes multiple ribosomal proteins from both the 30S and 50S subunits, but also factors involved in translation initiation, elongation, and termination, aminoacyl-tRNA charging, tRNA maturation and modification, rRNA modification, peptidyl-tRNA recycling, and RNA turnover. Thus, the translation-related signal is not limited to isolated ribosomal genes but extends across several layers of the gene-expression machinery. The identification of CosR-binding sites at rRNA and tRNA promoters adds an important dimension to the regulon. Direct regulation of rRNA promoters is a major mechanism by which bacteria couple ribosome production, nutrient availability, and growth rate (23–25). This organization is consistent with the essential role of CosR and raises the possibility that one of its major functions is to tune the translational capacity and growth rate of *C. jejuni* in response to environmental conditions.

**Figure 5.**
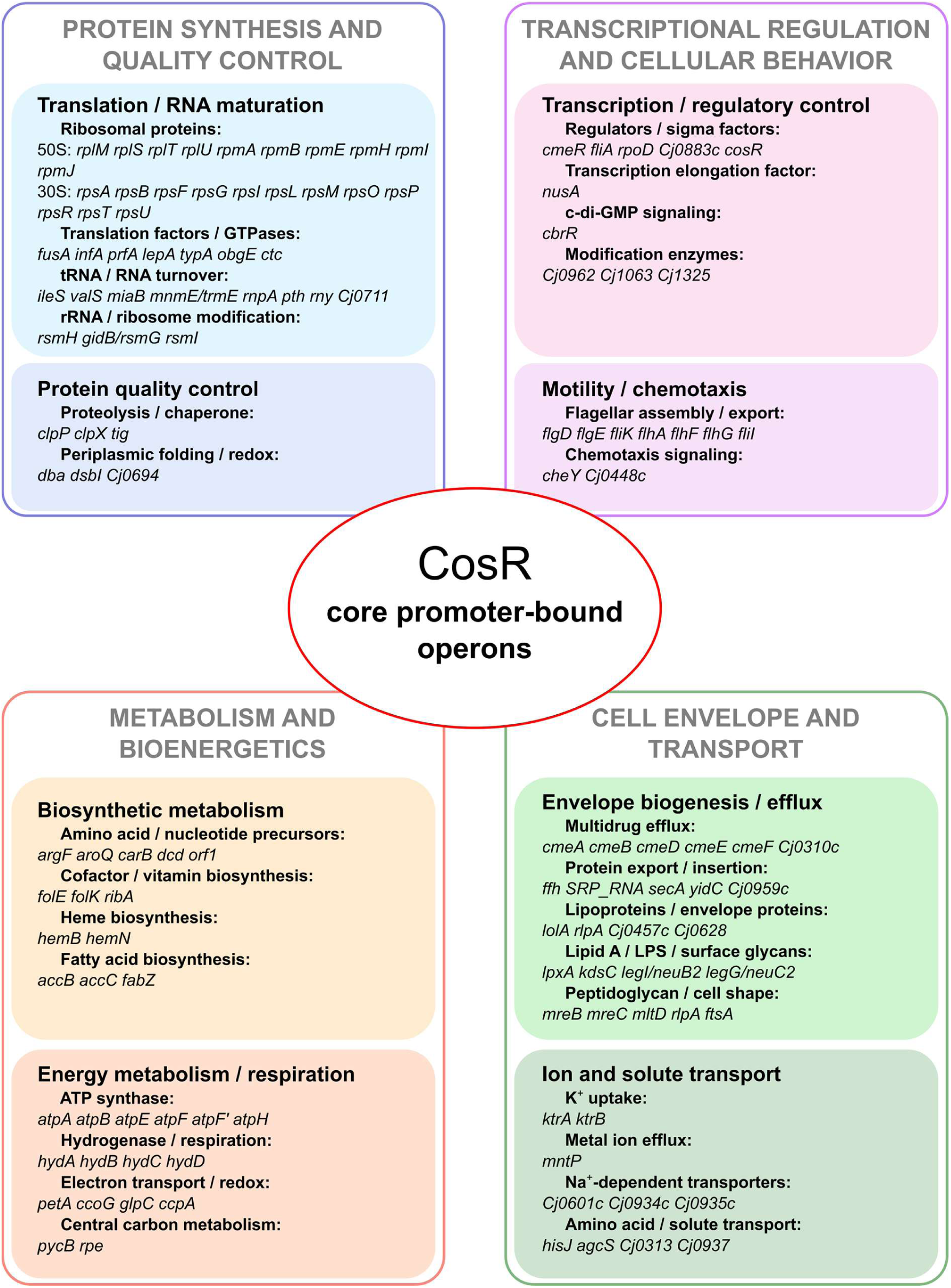
Functional organization of core promoter-bound CosR targets. Manual functional classification of genes belonging to core promoter-bound CosR-associated operons in *C. jejuni*. The eight functional categories were visually organized into four broader cellular domains. Only regulatory units associated with the stringent core ChIP-seq subset were included. Genes encoding poorly characterized or hypothetical proteins were omitted for clarity. Functional groups were manually assigned based on gene annotations and predicted biological functions, independently of COG categories used for enrichment analysis.

Beyond translation, the core promoter-bound regulon includes genes involved in transcriptional regulation, motility and chemotaxis, energy metabolism, respiration, membrane biogenesis, efflux, protein quality control, ion homeostasis, and biosynthetic pathways (Fig. 5). These categories are biologically coherent with the lifestyle of *C. jejuni*, which must rapidly adapt to changing environments during transmission, host colonization, and exposure to stress (4, 26). The additional presence of targets linked to ATP synthase, hydrogenase, electron transport, protein export, membrane insertion, flagellar assembly and efflux suggests that CosR coordinates central growth functions with adaptive traits relevant to host colonization (26, 27). Taken together, these observations argue for CosR as a broad physiological regulator rather than a dedicated stress-response factor.

The positional distribution of CosR-binding sites relative to the TSS shows a functionally structured pattern. Promoter-overlapping sites are associated with translation-related genes (COG J), whereas upstream sites are enriched in transcription and motility functions (COG K and COG N). This positional dichotomy suggests that CosR could act through distinct regulatory modes depending on promoter context: DNA binding close to the TSS could directly influence RNA polymerase recruitment, initiation or promoter clearance at ribosomal and translation-factor genes, while binding to genomic sites upstream of the core promoter may modulate RNA polymerase accessibility at motility and regulatory loci. Although the functional impact of these positional differences still needs to be experimentally investigated, the non-random distribution of binding sites suggests that CosR does not regulate all targets through a single, uniform mechanism. However, summit-based positional classes should be viewed as a genome-wide descriptive framework rather than as complete promoter-architecture models, since DNase I footprinting revealed multipartite binding patterns at several loci, with protected regions sometimes positioned differently from the ChIP-seq summit.

This view also helps reinterpret previous CosR studies. Earlier work linked CosR to oxidative-stress defense, multidrug efflux and additional physiological processes, mainly through antisense PNA-mediated knockdown or CosR overexpression (11, 16, 17). These approaches were important in establishing CosR as a global regulator, but they are difficult to interpret mechanistically because partial depletion of an essential protein can cause broad secondary effects on growth, metabolism, translation and stress physiology. Consistent with this limitation, the overlap between previously deregulated genes and promoter-bound targets identified here was limited. Among genes upregulated after PNA-mediated CosR depletion, only a small subset corresponded to operons associated with CosR-bound promoters in our dataset, including the *cj0040-fliK-flgD-flgE* and *cmeA-cmeB* regions. Conversely, very few downregulated genes overlapped with promoter-bound CosR targets, with *cj0776c-valS* representing one such case. This limited overlap suggests that earlier PNA-based regulons likely include both direct CosR targets and indirect transcriptional changes resulting from partial depletion of an essential regulator, with possible contributions from PNA toxicity and off-target effects.

The *cmeR-cmeABC* locus provides a useful benchmark for integrating our ChIP-seq dataset with previous CosR studies. Earlier work showed that CosR binds the *cmeABC* promoter and contributes, together with CmeR, to repression of the *cmeABC* multidrug efflux operon (15, 16). In our dataset, the strongest CosR signal in this region mapped to a previously undescribed site at the bidirectional promoter controlling *cj0369c-cmeR* and the divergently transcribed *rpsU* gene (Fig. S4). Manual inspection also revealed a weak, subthreshold CosR-enrichment signal upstream of *cmeABC*, consistent with the previously characterized *PcmeA*. Thus, our data refine rather than contradict the previous model: CosR may affect *cmeABC* both directly, through low-level binding at *PcmeA*, and indirectly, through regulation of the local repressor CmeR. The weak ChIP-seq signal at *PcmeA* may reflect the native repressed state of this promoter, where CmeR occupancy and limited promoter activity could reduce apparent CosR recovery. This supports direct CosR involvement in the *cmeR-cmeABC* module while extending the model to include regulation of the efflux operon and its local repressor.

In contrast, several oxidative-stress genes previously proposed as direct CosR targets were not recovered as robust promoter-bound targets in our ChIP-seq dataset. The *ahpC* and *luxS* promoter regions showed only weak, subthreshold enrichment in the unfiltered peak set (Supplementary Table S3), whereas those of *dps*, *sodB*, *katA*, and *perR* were not supported as high-confidence CosR-bound promoters under our conditions (11, 16, 17). These discrepancies do not exclude low-affinity, strain-specific or condition-dependent binding, particularly for sites detected by footprinting in previous studies. However, they argue against a model in which CosR primarily acts as a dedicated direct regulator of canonical oxidative-stress defense genes. Instead, the strongest reproducible signal in our data points to regulation of core physiological functions, especially the gene-expression machinery.

The DNA-binding properties defined here are compatible with this broader regulatory role. De novo motif analysis identified a ChIP-derived bipartite sequence signature, and motif analysis of footprint-protected regions recovered a statistically similar motif. Both signatures converge on two TTAA-like elements separated by an A/T-rich spacer, although individual nucleotide preferences differ between the ChIP-derived and footprint-derived logos. This is not unexpected, because ChIP-seq captures enriched summit-centered regions in vivo, whereas footprinting defines local protected DNA segments in vitro. Together, these data suggest that CosR recognizes a conserved bipartite sequence logic embedded in structurally diverse promoters, rather than a rigid invariant linear consensus.

This flexible recognition model fits well with the structural properties of CosR. Structural studies showed that apo-CosR forms a symmetric dimer, whereas DNA binding induces an asymmetric arrangement in which the two protomers adopt different conformations (13). The winged helix-turn-helix domain contacts the major groove, while the wing contributes to minor-groove interaction and stabilization. Such conformational plasticity could allow CosR to accommodate variable spacing, partial half-site conservation, and promoter-dependent DNA shape features. This may explain why CosR-protected regions differ among promoters despite sharing an AT-rich recognition logic. It is also consistent with the similarity between CosR and the *Helicobacter pylori* HP1043/HsrA regulator, whose DNA-binding domain is highly conserved and whose regulon is also enriched in central physiological functions (14, 28). Thus, CosR and HP1043 may represent related regulatory solutions in Campylobacteria, linking essential promoter recognition to control of core cellular processes.

Direct binding to the *cosR* promoter further supports an autoregulatory loop. Autoregulation is common among transcription factors that control essential or growth-linked processes, because their intracellular concentration must be tightly tuned. In this context, CosR autoregulation may contribute to maintaining appropriate regulator levels while allowing rapid adjustment under changing physiological conditions. Direct binding at rRNA and tRNA promoters further strengthens the view that CosR acts at the level of the gene-expression apparatus itself.

The oxidative-stress experiments indicate that CosR is condition responsive, but also reveal that CosR occupancy and transcriptional output are not linearly coupled. A short H₂O₂ treatment increased CosR binding at a subset of promoters, including *atpF′*, *cbrR*, *flaC*, and *cosR*, but reduced binding at translation-associated promoters such as *rpsF*, *rplS*, *rpmJ*, and *tRNASer*. The redox footprinting experiments support a direct effect of oxidation on CosR-DNA interaction, since H₂O₂ strongly reduced CosR-dependent protection at the *rplS* promoter and similarly impaired protection at *atpE*. This is consistent with previous evidence that oxidation impairs CosR binding to the *cmeABC* promoter and that substitution of the only cysteine residue of CosR, C218, renders the protein insensitive to oxidation (15). Thus, C218-dependent redox modulation provides a plausible biochemical mechanism for reduced CosR-DNA binding at multiple promoters.

The increased ChIP enrichment observed at other promoters should therefore not necessarily be interpreted as an opposite intrinsic response of CosR to oxidation. In vivo occupancy also reflects promoter context, DNA topology, RNA polymerase occupancy, and additional stress-responsive factors. One parsimonious interpretation is that oxidation reduces CosR binding at susceptible sites, including translation-associated promoters, while the apparent increase at other promoters reflects changes in the in vivo regulatory context. At the transcriptional level, most tested genes were downregulated after H₂O₂ treatment regardless of whether CosR occupancy increased or decreased at the corresponding promoter. This uncoupling suggests that oxidative stress remodels CosR binding, but that the final output also depends on promoter architecture and additional regulatory inputs. This is consistent with the complex oxidative-stress regulatory network of *C. jejuni*, in which peroxide response, iron homeostasis and redox-responsive regulators are functionally interconnected (8–10). Therefore, while direct oxidation of CosR is likely part of the response, the mixed in vivo occupancy pattern indicates that promoter-context effects and additional regulators remain to be dissected.

Taken together, our data redefine CosR as an essential promoter-binding regulator that directly controls central cellular processes in *C. jejuni*. Its core promoter-bound regulon is dominated by translation and RNA maturation, but also includes regulatory, metabolic, respiratory, motility, envelope, efflux and protein-quality-control functions. CosR recognizes bipartite TTAA-like sequence features within heterogeneous promoter architectures, binds its own promoter, and rapidly remodels promoter occupancy under oxidative stress. These findings position CosR as a condition-responsive regulator linking promoter recognition, core physiology and stress-associated transcriptional reprogramming.

## Author contributions

Elena Chiti: Investigation, Writing – original draft, Writing – review & editing. Tobia Odorici: Investigation, Software, Writing – review & editing. Mateusz Noszka: Investigation, Writing – review & editing. Jakub Muraszko: Investigation, Writing – review & editing. Annamaria Zannoni: Investigation, Writing – review & editing. Anna Zawilak-Pawlik: Resources, Supervision, Writing – review & editing. Andrea Vannini: Conceptualization, Data curation, Investigation, Project administration, Resources, Software, Supervision, Visualization, Writing – original draft, Writing – review & editing. Davide Roncarati: Conceptualization, Investigation, Project administration, Resources, Supervision, Visualization, Writing – original draft, Writing – review & editing.

## Conflict of interest

The authors declare no conflict of interest.

## Funding

This work was supported by institutional research funds from Alma Mater Studiorum – University of Bologna; no dedicated project grant was used.

